# Outbreak of *Dermatophilus congolensis* skin infection among contact sport practitioners, Norway, summer 2025

**DOI:** 10.64898/2026.07.24.740579

**Authors:** Kristine Pape, Sveinung Javnes, Christina G. Aas, Eli O. Sagvik, Kjersti W. Larsen, Solveig Jore

**Affiliations:** Municipality of Trondheim, Department of Infectious Disease Control, Trondheim, Norway; Norwegian University of Science and Technology, Trondheim, Norway; St. Olavs Hospital, Trondheim University Hospital, Department of Medical Microbiology, Trondheim, Norway; Norwegian Institute of Public Health, Oslo. Norway

**Keywords:** Dermatophilus, Dermatophilosis, Disease Outbreaks, Communicable Disease Transmission, Contact Tracing, Contact Sports, Skin diseases, Bacterial, Molecular Epidemiology, Whole Genome Sequencing, Public Health Surveillance, Norway

## Abstract

*Dermatophilus congolensis* is a zoonotic gram-positive bacterium causing dermatophilosis, a skin infection primarily affecting animals and uncommonly reported in humans. Although human infections have traditionally been associated with animal contact, recent reports suggest alternative transmission pathways. We investigated an outbreak among contact sport athletes in Trondheim, Norway, during July to September 2025, using structured interviews and whole-genome sequencing. Nine confirmed cases were identified. Symptoms were mild and consisted mainly of pustular lesions affecting the face, arms, back, and chest. None of the cases reported animal contact. Genomic analysis confirmed species identity and showed that outbreak isolates were highly related, differing by only 0-7 core-genome single nucleotide polymorphisms. The Trondheim isolates clustered with strains from recent outbreaks in Spain and France, suggesting international dissemination of a shared clone. These findings identify contact sports as a potential setting for transmission of human dermatophilosis and highlight the value of genomic surveillance for outbreak investigation.

## Background

*Dermatophilus congolensis* is a gram-positive bacterium belonging to the *Dermatophiliaceae* family within the order *Actinomycetales. D. congolensis* is primarily an animal pathogen and an opportunistic zoonotic pathogen in humans, causing dermatophilosis. Transmission occurs via direct contact with infected animals or indirectly through contaminated environments or materials, with highest prevalence in warm, humid tropical and subtropical regions *(1, 2)*. In addition to climatic factors, a broken skin barrier due to wounds, insect bites or microtrauma, seem essential for an infection to establish upon exposure *(1)*.

Human infection with *D. congolensis* has been documented as sporadic cases or small outbreaks over the last 65 years *(1, 3-15)*. Most described cases have been associated with confirmed or suspected direct or indirect contact with infected animals, with a few exceptions. Until recently, European cases have mainly been restricted to travelers or workers returning from tropical regions, and with suspected animal exposures *(8-13)*.

However, in June 2026, outbreaks of human dermatophilosis among men who have sex with men (MSM) were reported from Spain and France *(16, 17)*, with additional related cases subsequently identified in Germany and Sweden *(18, 19)*. These observations suggest that transmission may occur in the absence of animal contact.

Clinically, human dermatophilosis typically presents as localized skin lesions such as pustules, papules and dermatitis, although atypical manifestations have also been described *(20-22)*. The incubation period appears to be short, with lesions usually occurring within a few days of exposure *(4, 19)*. The infection is usually self-limiting, but may take a chronic course in some *(23)*. Dermatophilosis may be underrecognized, as infections are often mild and clinically indistinguishable from more common skin conditions *(9)*.

Here, we report a cluster of *D. congolensis* infections occurring in a grappling-based contact sport environment in Norway, representing a different epidemiological setting characterized by close skin-to-skin contact and repeated minor skin trauma. To our knowledge, this is the first report of an outbreak among contact sport practitioners due to *D. congolensis* and the northernmost outbreak of human dermatophilosis described to date.

### Outbreak detection

On August 6, 2025, the Municipal Medical Officer in Trondheim was notified by a general practitioner of a laboratory-confirmed *D. congolensis* skin infection in a male athlete. A second laboratory-confirmed case from the same grappling-based sports club was identified shortly thereafter, raising suspicion of an outbreak. Because *D. congolensis* is rarely reported in Norway, local and national public health authorities initiated an outbreak investigation. Through active case finding, seven additional cases were identified during the following six weeks. This report describes the epidemiological investigation, microbiological findings, and control measures implemented to end the outbreak.

## Methods

### Outbreak setting

The outbreak occurred in Trondheim, a mid-sized urban municipality situated in central Norway (lt 63.4 N) with about 218,000 inhabitants. Summer in Trondheim is moderately wet, and average daytime temperatures typically range from 17-20 °C in July and August. July 2025 was, however, noticeably warmer with an average mid-temperature 3,9 °C above normal. During a heatwave between July 12 and July 24 daytime temperatures reached around 30 °C, far exceeding typical seasonal levels *(24)*.

### Case definition

During the outbreak we defined a case as:

- **Confirmed**: A person training at the contact sports club with a laboratory-confirmed infection with *D. congolensis*, with symptom onset or bacterial wound sampling date after 20 July 2025
- **Probable**: A person living in Norway with a laboratory-confirmed infection with *D. congolensis* with symptom onset or bacterial wound sampling date after 20 July 2025.

The retrospective case definition included all *D. congolensis* with single linkage clustering of ≤ 10 core-genome single nucleotide polymorphisms (SNPs) following whole genome sequencing (WGS).

### Epidemiological investigation

Other microbiological laboratories were contacted regarding recent positive cultures of *D. congolensis*. In addition, a systematic search was performed in the laboratory data information system of St. Olavs Hospital to identify all human cases of *Dermatophilus spp*. in Trondheim in the past 5 years.

A structured interview guide was developed based on knowledge from literature and the current outbreak. Topics included personal information, clinical presentation and course, health care contacts including treatment/interventions, information on other known cases and possible causes. Targeted questions addressed close contacts, training habits, travel within 3 months prior to symptoms, and potential exposures to training facilities, animals (in particular horses and farm animals), and fresh-water swimming (see Appendix for Interview guide).

The nine confirmed cases agreed to participate in the interview and provided written consent (see Appendix for Consent form). Two municipal team members conducted structured 30-minute telephone interviews in October 2025. For the probable case, no informed consent for interview was achieved.

### Microbiological investigation

Skin swab samples from 10 athletes with pustulosis (n=10) were submitted on Amies liquid medium (Eswab, Copan), inoculated on Columbia blood agar with 5% sheep blood, MacConkey agar and chocolate blood agar and incubated at 37°C in 5% CO_2_ for 24-48h.

Identification was performed by phenotypical appearance on blood agar and in Gram stain in addition to matrix-assisted laser-desorption/ionization time-of-flight mass spectrometry (MALDI-TOF MS) using the MALDI Biotyper® Sirius system (Bruker Daltonics) and the MALDI Biotyper Compass Library (V13.0.0.2_11897-12438) *(25, 26)*. Both direct plating and on-plate extraction with formic acid were performed. A score value ≥ 2.0 was regarded as valid for identification to species level and a score value 1.7 - < 2.0 was regarded as valid for identification to genus level according to the manufacturer’s recommendations. Identification was confirmed with whole genome sequencing for 10 of the isolates. One of the isolates had not been frozen and was only subject to conventional identification, but not to susceptibility testing or sequencing.

Antimicrobial susceptibility testing was performed by broth microdilution using Mueller Hinton broth and Sensititre™ EUSTAPH supplied with benzylpenicillin from Sensititre™ STP6F (Thermo Scientific). Minimum inhibitory concetration (MIC) was interpreted according to EUCAST recommendations for aerobic gram-positive organisms *(27)*.

Genomic DNA was extracted from overnight cultures grown in tryptic soy broth at 37 °C. Cell pellets were incubated for 2–3 h at 37 °C in lysis buffer (10 mM EDTA, 100 mM NaCl, 0.5% Triton X-100, 0.5% Tween-20) containing mutanolysin, lysozyme, and proteinase K. After incubation at 65 °C, RNAse A (100 mg/mL) was added, followed by automated extraction using EZ1&2 DNA Tissue kit on an EZ1 Advanced XL extractor.

Ten isolates were whole genome sequenced using Illumina technology. Briefly, sequencing libraries were prepared with the DNA prep kit and sequenced on a MiSeq instrument (Illumina) using the MiSeq V3 reagent kit (2×300 bp). Four isolates were additionally sequenced using Oxford Nanopore technology, with the rapid sequencing kit (SQK-RBK114.24 V14) and a MinION Mk1b instrument (Oxford Nanopore Technologies). Basecalling was performed with Dorado v7.11.2 (sup@v5.2.0 model).

Quality control and trimming of Illumina data was performed using fastQC *(28)* and trimmomatic *(29)*. Taxonomic classification was performed with kraken2 *(30)* and the PlusPF database. Assembly was performed by Shovill *(31)* and annotation with Prokka *(32)*. Sequence data of cases is available from GenBank Bioproject PRJNA1483954. The core genome (1.52 Mbp) was defined using Roary *(33)* and a ML phylogeny inferred using Fasttree *(34)* with GTR model. Pairwise SNP distances were calculated using megaCC. Nanopore data were filtered using Filtlong v0.2.1 (--min_length 1000--keep_percent 90) *(35)*, assembled using Flye v2.9.4 *(36)* and polished using Illumina data with Pypolca v0.3.1 (--careful). Datasets for *D. congolensis* reference genomes and isolates from outbreaks in Spain *(17)* and France *(16)* were downloaded from ENA and processed as described above.

Potential antimicrobial resistance genes were detected using, AMRFinderplus, Abricate (VFDB and ResFinder databases) *(37)* as well as CARD RGI *(38)*. Illumina raw data were mapped against the tetZ gene of *D. congolensis* BTSK9 7 (NZ_JAAFON010000005).

## Results

### Epidemiological investigation

Between 29 July and 17 September 2025, 9 cases were confirmed, with symptom onset between 20 July and 14 September (Figure 1, upper section). In addition, one probable case was identified in south-eastern Norway 31 December 2025. The systematic laboratory data search from Trondheim identified one single report of *D. congolensis* from 2024 that was included in molecular analyses. No additional *Dermatophilus spp*. were identified in Trondheim during the past 5 years.

**Figure 1.**
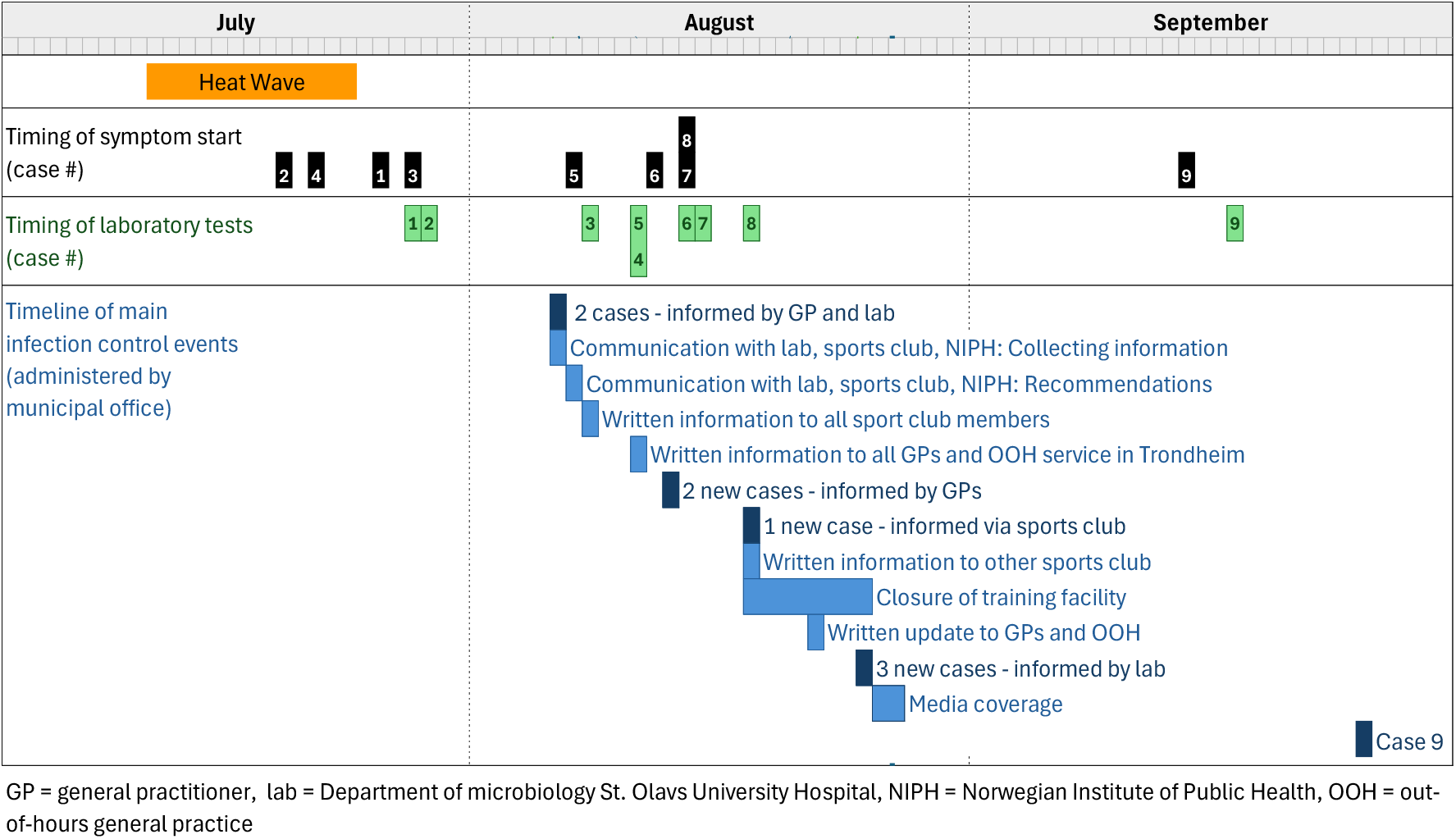
Visual overview of the outbreak, including timing of main events: symptom start (black), microbiological sampling (green) and infection control measures (blue).

All nine confirmed and interviewed cases were active practitioners of a grappling-based contact sport in the same club. Their median age was 40 years (range: 29–55), eight were male and all lived in stable relationships in a family setting, 8/9 with children in the household.

All interviewed cases described pustular or follicular skin lesions, initially involving the face (under eye, chin, forehead) or upper extremities (biceps, wrist). Lesions subsequently spread to additional body sites in 8/9 cases, most commonly the arms, face, chest and back. Only one case reported involvement of the abdomen and lower extremities. Symptoms were generally mild, with pruritus reported by six cases (6/9) and no systemic manifestations. Symptom duration ranged from 5–25 days (median: 20 days). The probable case had skin lesions in the face, scalp and knees upon contact with the out-of-hours general practitioner service in Oslo.

All interviewed cases had contacted their GP shortly after symptom onset. The four cases with symptom onset in July (cases #1-4), before an outbreak was suspected, all received various topical treatments (e.g., clindamycin liniment, bacitracin ointment) and two cases also received systemic antibiotics on suspicion of *Staphylococcus aureus* infection. The two cases initially receiving topical treatment alone reported limited effect and symptom progression. Once an outbreak was suspected all were recommended topical treatment with Chlorhexidine /Hibiscrub wash and prescription of systemic antibiotics. With systemic antibiotics, (phenoxymethylpenicillin (n=3), dicloxacillin (n=3), amoxicillin (n=1), doxycycline (n=1) and lymecycline (n=1)), improvement of symptoms was reported within 2-14 days.

All interviewed cases reported practicing contact sport regularly, usually at least three times weekly. The most common clothing was t-shirt/rash guard and shorts/spats/tights or Gi (uniform with pants, jacket and belt). Three cases had attended training sessions regularly during the whole month of July until symptom onset (cases #1, #3 and #8), while the rest had taken up practice after returning from vacation in mid-July (3/9, cases #2, #4 and #5) or the beginning of August (3/9, cases #6, #7 and #9). All cases had been attending training sessions during the period immediately before symptom onset. During interviews, cases also provided relevant information regarding the concept and practice of “open mat” during summer, meaning that anyone in the international community may show up and participate in the daily training sessions. Also, one case described the training environment during the heat wave period as “tropical”, with moist floor/mats.

None of the cases reported any symptoms in household members or other close contacts outside the sports club. None of the cases reported close contact with farm animals or horses during the three months prior to symptom onset, neither domestically nor abroad. Six of nine cases reported travel abroad within the preceding three months, all within Europe. Of these, four (4/9) had trained in clubs abroad (Sweden, Wales and/or Spain) of which two had attended the same sports club in Spain directly before returning home in the middle of July. They were not aware of other practitioners exhibiting symptoms at the Spanish club. Several cases had been swimming in the sea and in local lakes during the heat period in July. The probable case had recently returned home from contact sport practice in South-East Asia.

### Microbiological investigation

All samples but one grew pure cultures of typical adherent, white, wrinkled colonies with betahaemolysis (Figure 2A) and a characteristic gram stain displaying filamentous forms with transverse and longitudinal segmentation as well as coccoid forms (Figure 2B) within 2 days. One of the samples also recovered growth of methicillin-susceptible *S. aureus*.

**Figure 2.**
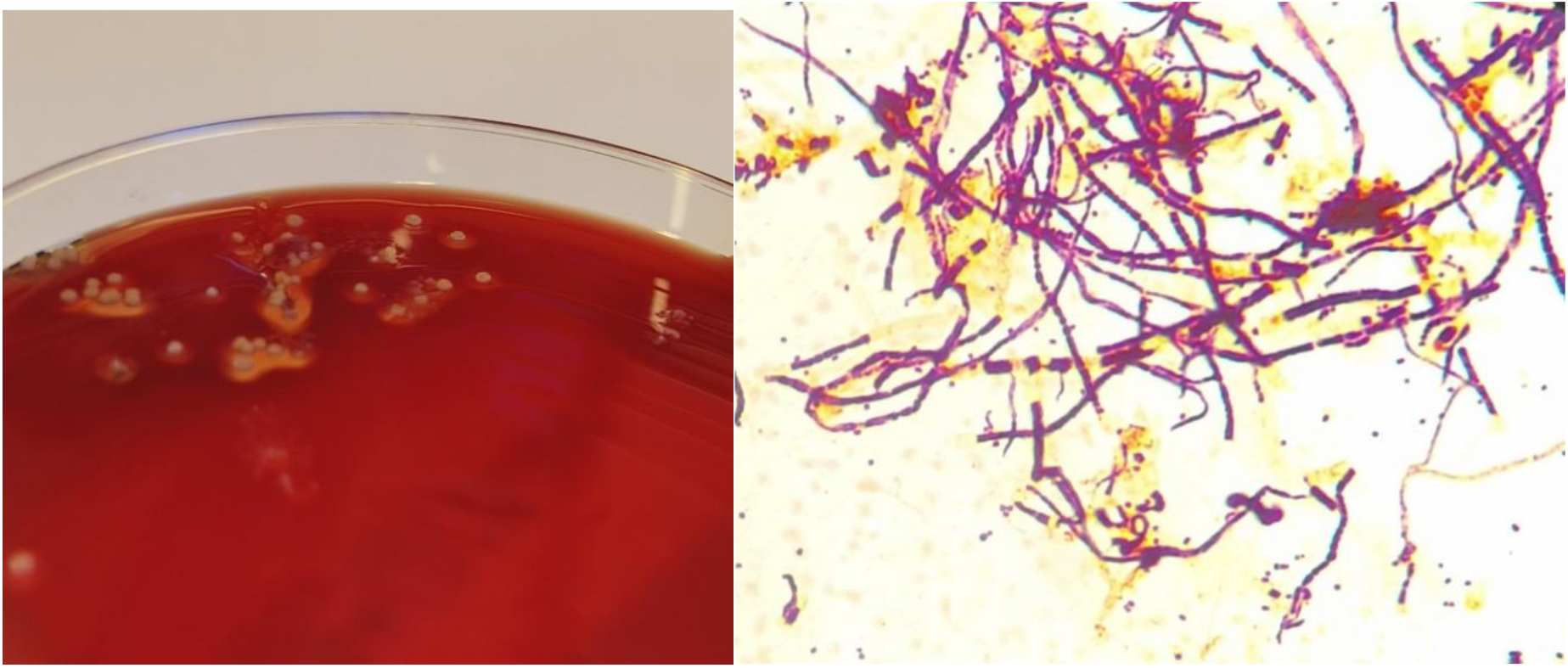
A.D. congolensis characteristic colonies B. Gram stain displaying filamentous and coccoid forms

MALDI-TOF MS analysis of all outbreak isolates yielded score values ranging from 1.47 to 1.66 with *D. congolensis* as the first suggested species in all cases. On-plate extraction yielded no added benefit for identification scores.

Nine isolates showed identical susceptibility profiles for clinically relevant antimicrobials: Tetracyclin MIC ≤ 0.5, benzylpenicillin MIC ≤ 0.06 mg/L, vancomycin MIC ≤ 0.5 mg/L, trimetoprime-sulfamethoxazol MIC ≤ 1 mg/L, clindamycin MIC ≤ 0.25 mg/L, moxifloxacin MIC ≤ 0.25 mg/L and linezolid MIC ≤ 2 mg/L. These would be regarded as agents that could be considered for use in therapy according to Eucast recommendations. In addition, all isolates showed erythromycin MIC ≤ 0.25 mg/L, fucidic acid MIC ≤ 0.5 mg/L and mupirocin MIC > 256 mg/L. For these agents there are no recommendations for interpretation.

Whole genome sequencing confirmed the initial species identity of the 10 sequenced isolates, displaying 82.4-93.5 % match to *D. congolensis*. Core genome phylogeny revealed close genetic relatedness between 9 of these isolates, including the probable case, with pairwise SNP distances ranging from 0-7 (Figure 3 and Figure 4 (Appendix)), in support of the epidemiological data. The single isolate from 2024 clustered separately, with more than 8,900 SNP differences from the outbreak cluster and was thus regarded as unrelated.

**Figure 3.**
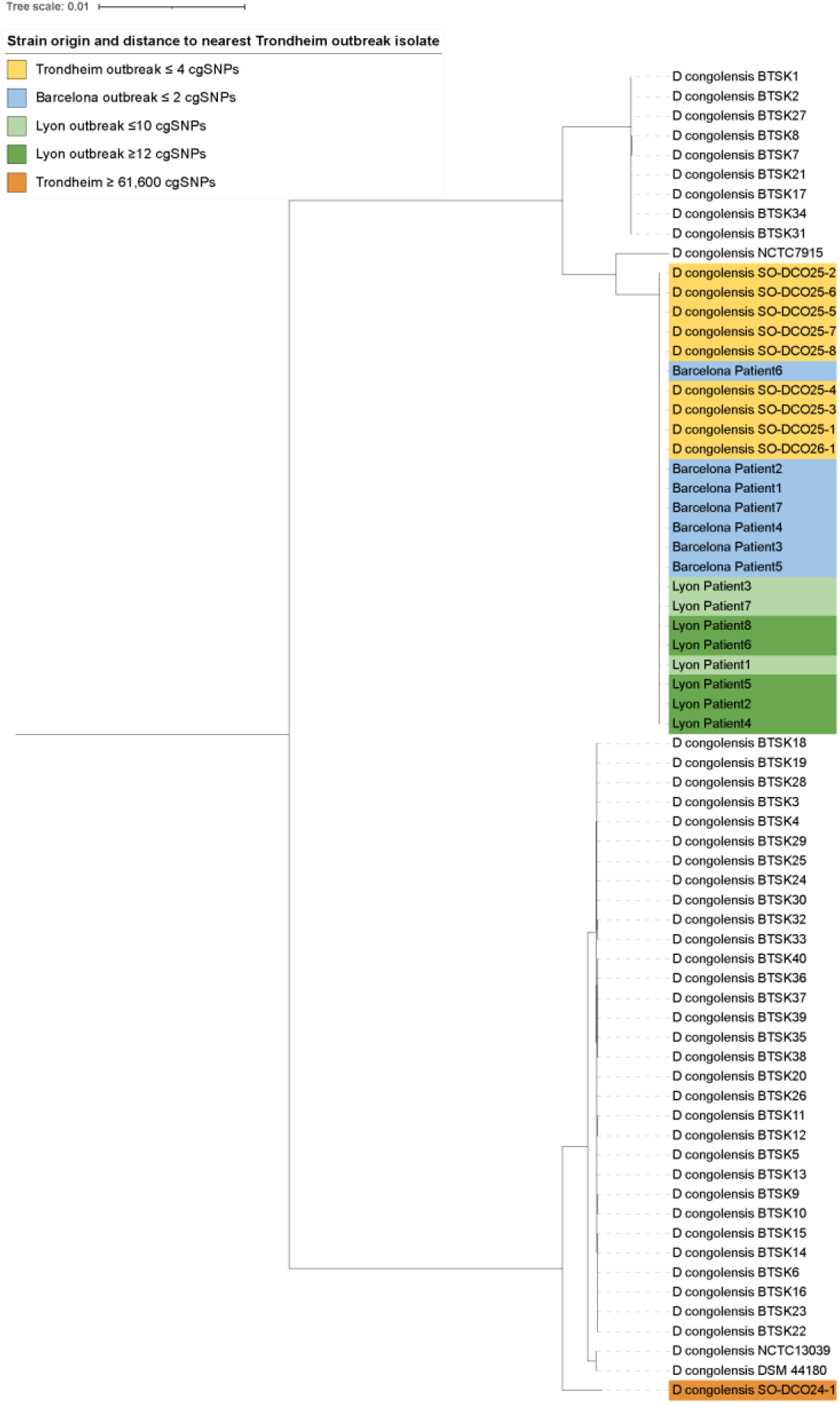
Core genome phylogeny (1.52 Mbp) of D. congolensis isolates and reference genomes. Isolates from outbreaks in Trondheim, Barcelona and Lyon are highlighted, with SNP distance to the nearest Trondheim outbreak isolate provided in the legend.

Combining nanopore and Illumina data produced a complete genome for the outbreak strain, consisting of a single circular 2.63 Mbp chromosome with a GC of 59.0% encoding 2,413 genes. The most closely related reference genome to the outbreak strain was *D. congolensis* NCTC7915, previously shown to belong to a separate cluster of *S. congolensis (39)*. Using multiple tools and databases, we were unable to detect any significant similarity to known antibiotic resistance genes (including the *tetZ* gene of *D. congolensis* BTSK9).

Notably, the outbreak strains clustered together with strains from the recently reported outbreaks in Barcelona and Lyon (Figure 3 and Figure 4 (Appendix)) with pairwise core genome SNP distances ranging from 1-7 to the Spanish isolates and 3-52 to the French isolates, suggesting a potential epidemiological link.

### Control measures

Control measures were coordinated and implemented by the municipal office in collaboration with local and national expertise. An overview and timeline are presented in the lower part of Figure 1. Infection control measures included recommendations to the leader and individual practitioners at the sports club, dissemination of information to local general practitioners and thorough cleaning of the training facilities with soap and water, including training mats and contact points. In addition, it was decided to close the training facility for one week from 18 August. A detailed list of specific measures, recommendations and timeline is provided in the Appendix (Control measures).

## Discussion

We report an outbreak of *D. congolensis* in Trondheim, Norway, comprising nine confirmed cases between 29 July and 17 September 2025, with an additional genetically linked Norwegian case from December 2025. All confirmed cases occurred among practitioners of a grappling-based contact sport within the same club. To our knowledge, this represents the first reported outbreak of dermatophilosis associated with contact sports in Europe, and possibly the first described in this setting globally. Epidemiological investigation did not identify animal exposure in any of the cases, and no infections were observed among partners, children, or other contacts outside the sports club.

Until recently, *D. congolensis* has been regarded primarily as an animal pathogen, occasionally causing zoonotic infections in humans. Transmission from infected animals has been suggested to occur through both direct contact and indirect exposure via contaminated materials or environments *(2)*. The present outbreak, together with recently reported European outbreaks among men who have sex with men *(16-19)*, suggests that transmission can occur in the absence of animal contact and beyond traditional tropical and subtropical settings. The Trondheim cluster represents the northernmost outbreak described to date and demonstrates that human dermatophilosis is not restricted to MSM-associated transmission networks.

In our cluster, lesion distribution differed from that reported in MSM outbreaks, with initial involvement of areas commonly exposed during training such as the arms and face. While compatible with direct transmission, this pattern, together with spread to the back, suggests that contaminated mats, clothes, or equipment may also have contributed. The absence of secondary transmission in households indicates limited transmissibility and suggests that additional factors such as skin trauma, repeated wetting, and local heat and humidity may be important. Direct and indirect transmission pathways may therefore operate in parallel, although their relative contribution remains uncertain. Environmental sampling of mats and other high-contact surfaces could have been valuable but was not performed during the outbreak.

Genetic analyses showed that the Trondheim isolates clustered closely with isolates from Barcelona and showed relatedness to isolates from France. An additional case (our probable case) from south-eastern Norway associated with travel to South-East Asia also clustered with the same group. Although directionality cannot be inferred, these findings support the hypothesis of international spread of a single strain or clone, likely introduced during summer 2025 and subsequently disseminated through interconnected training environments.

Contact sport settings are well known for facilitating transmission of skin pathogens *(40, 41)*. Grappling activities involve prolonged skin-to-skin contact, repeated friction, and minor skin injuries that disrupt the skin barrier and may facilitate entry of infectious agents, including *D. congolensis* zoospores. The transmission pattern observed in this outbreak resembles that described for other infections associated with contact sports, including *S. aureus*, herpes gladiatorum, and dermatophyte infections *(42)*. Although direct contact is likely important, contaminated mats and equipment may also contribute *(43)*. Modern training practices may further facilitate dissemination across borders. Open-mat sessions and international seminars and competitions involve close contact between individuals from different regions and countries. In the present outbreak, the infection was most likely introduced by practitioners who trained at a club in Spain during a family vacation. Together with participation in international competitions and a genetically related case associated with travel and contact sport practice in South-East Asia, these findings support the hypothesis that international contact networks may facilitate the spread of uncommon pathogens across geographic regions.

Environmental conditions may also have contributed to this outbreak. The first cases occurred during an unusually warm period in Trondheim, during which indoor training conditions were described as exceptionally hot and humid. Warmth and moisture are well-known risk factors for dermatophilosis because prolonged skin hydration and minor skin trauma facilitate invasion by *D. congolensis*. Although climate cannot be implicated directly, our observations are consistent with the understanding that dermatophilosis is a climate-sensitive infection. As temperatures increase and extreme weather events become more frequent, environmental conditions favorable for infection may become more common, potentially influencing the occurrence and geographic distribution of dermatophilosis and other skin-associated infectious diseases *(44)*.

*D. congolensis* is an uncommon finding in clinical human samples in Norway. Although no national registry for human cases exists, only one genetically unrelated isolate was identified in Trondheim in the five years preceding this outbreak. Given that the organism can be cultured using standard methods, this suggests that infections are uncommon in Norway. In this outbreak, MALDI-TOF MS yielded low identification scores despite repeated attempts, likely reflecting a limited representation of *D. congolensis* in the database. This is consistent with known limitations of MALDI-TOF MS for other *Actinobacteria*, where identification may depend on database completeness and may require adjusted score thresholds or complementary methods *(26)*. In contrast, the genetically unrelated isolate from 2024 yielded a higher identification score, suggesting better representation of that strain in the database. If more spectra from *D*.*congolensis* strains of different genetic backgrounds are added to the MALDI-TOF MS database, this issue will likely improve.

Antimicrobial susceptibility testing showed low MIC values for commonly used agents, including benzylpenicillin, doxycycline, clindamycin, and trimethoprim-sulfamethoxazole, and no known antimicrobial resistance genes were identified. However, no standardized testing methodology or clinical breakpoints exist for *Dermatophilus* spp., and susceptibility results should therefore be interpreted cautiously.

This study has limitations. Additional people with dermatophilosis may not have been detected, although no further local cases were identified after the last confirmed case. The number of cases was small and the presentation of details regarding the timeline and exposures is limited to prevent person identification. Also, interviews were performed retrospectively, introducing potential information bias, especially regarding timing of exposures. The structured interviews were also elaborated based on available knowledge at the time, and information regarding sexual activities was not collected. Despite these limitations, the combination of epidemiologic data and whole-genome sequencing provides strong evidence for a common outbreak strain.

The outbreak was successfully controlled through early case identification, antimicrobial treatment, enhanced surveillance, and hygiene measures targeting both practitioners and the training environment. Close cooperation with the sports club and high awareness among participants were essential for limiting further spread. In conclusion, this outbreak demonstrates that *D. congolensis* can cause clusters of infection in contact sport settings when environmental and behavioral conditions are favorable. The findings support a model in which skin trauma, moisture, and repeated close contact are key determinants for transmission and highlight the potential for international dissemination within connected training communities. Finally, this study illustrates the value of genomic surveillance for outbreak investigation.

## References

1. Zaria LT. Dermatophilus congolensis infection (Dermatophilosis) in animals and man! An update. Comp Immunol Microbiol Infect Dis. 1993 Jul;16(3):179–222.

2. Banwo OG, Akinsulie OC, Adesola RO, Jeremiah OT. Dermatophilosis: Current Advances and Future Directions. Acta Microbiologica Hellenica. 2025;70(4):40.

3. Burd EM, Juzych LA, Rudrik JT, Habib F. Pustular dermatitis caused by Dermatophilus congolensis. J Clin Microbiol. 2007 May;45(5):1655–8.

4. Dean DJ, Gordon MA, Severinghaus CW, Kroll E, Reilly JR. Streptothricosis: a new zoonotic disease. New York state journal of medicine. 1961;61.

5. Dusch H, Huszar A, Nicolet J, Graevenitz Av, Collins MD. Characterisation of an unusual human isolate of Dermatophilus congolensis. Medical Microbiology Letters. 1994;3(1):36–41.

6. Erickson EL. Dermatophilus congolensis infection in man. Cutis. 1975;16(1):83–4.

7. Kaminski GW, Suter, II. Human infection with Dermatophilus congolensis. Med J Aust. 1976 Mar 27;1(13):443–7.

8. Londero AT, Ramos CD, Souza LP. Human dermatophilosis, its occurrence in Brazil. Mykosen. 1974 May 1;17(5):111–3.

9. Towersey L, Martins Ede C, Londero AT, Hay RJ, Soares Filho PJ, Takiya CM, et al. Dermatophilus congolensis human infection. J Am Acad Dermatol. 1993 Aug;29(2 Pt 2):351–4.

10. Alejo-Cancho I, Bosch J, Vergara A, Mascaro JM, Marco F, Vila J. Dermatitis by Dermatophilus congolensis. Clin Microbiol Infect. 2015 Sep;21(9):e73–4.

11. Amor A, Enriquez A, Corcuera MT, Toro C, Herrero D, Baquero M. Is infection by Dermatophilus congolensis underdiagnosed? J Clin Microbiol. 2011 Jan;49(1):449– 51.

12. Aubin GG, Guillouzouic A, Chamoux C, Lepelletier D, Barbarot S, Corvec S. Two family members with skin infection due to Dermatophilus congolensis: a case report and literature review. Eur J Dermatol. 2016 Dec 1;26(6):621–2.

13. de Lorenzi C, Quenan S, Fontao L. Dermatophilus congolensis dermatitis in a traveller from Thailand. Journal of Travel Medicine. 2021;28(6).

14. Porras MI, Canueto J, Ferreira L, Garcia MI. Human dermatophilosis. First description in Spain and diagnosis by matrix-assisted laser desorption ionization time-of-flight mass spectrometry (MALDI-TOF). Enferm Infecc Microbiol Clin. 2010 Dec;28(10):747–8.

15. Pospisil L, Skalka B, Bucek J, Moster M. [The first isolation of Dermatophilus congolensis van Saceghem 1913 in Czechoslovakia]. Cesk Epidemiol Mikrobiol Imunol. 1992 Oct;41(5):258–67.

16. Degreze M, Durupt F, Ibranosyan M, Maucotel AL, Lapendry A, Gouillon L, et al. Suspected Sexual Transmission of Dermatophilosis among Men Who Have Sex with Men, Lyon and Paris, France, 2025-2026. Emerg Infect Dis. 2026 Jun;32(6):959–63.

17. Descalzo V, Moreno-Mingorance A, Alvarez-Lopez P, Salmeron P, Garcia-Perez JN, Pericas-Cladera FP, et al. Suspected Sexual Transmission of Dermatophilosis among Men Who Have Sex with Men, Barcelona, Spain, 2025-2026. Emerg Infect Dis. 2026 Jun;32(6):964–9.

18. European Centre for Disease Prevention and Control. Rapid risk assessment – Clusters of dermatophilosis in five EU/EEA countries in 2025–2026. 17 June 2026. ECDC: Stockholm; 2026.

19. Filen F, Loo E, Haij Bhattarai K. Dermatophilosis among men who have sex with men, Stockholm, Sweden, March to June, 2026. Euro Surveill. 2026 Jun;31(25).

20. Bunker ML, Chewning L, Wang SE, Gordon MA. Dermatophilus congolensis and “hairy” leukoplakia. Am J Clin Pathol. 1988 May;89(5):683–7.

21. Gillum RL, Qadri SM, Al-Ahdal MN, Connor DH, Strano AJ. Pitted keratolysis: a manifestation of human dermatophilosis. Dermatologica. 1988;177(5):305–8.

22. Harman M, Sekin S, Akdeniz S. Human dermatophilosis mimicking ringworm. Br J Dermatol. 2001 Jul;145(1):170–1.

23. Albrecht R, Horowitz S, Gilbert E, Hong R, Richard J, Connor DH. Dermatophilus congolensis chronic nodular disease in man. Pediatrics. 1974 Jun;53(6):907–12.

24. Seklima - Observations and weather statistics [internet]. [cited 2026 Jun 20]; https://seklima.met.no/observations/

25. Cuenod A, Foucault F, Pfluger V, Egli A. Factors Associated With MALDI-TOF Mass Spectral Quality of Species Identification in Clinical Routine Diagnostics. Front Cell Infect Microbiol. 2021;11:646–648.

26. Schulthess B, Bloemberg GV, Zbinden R, Bottger EC, Hombach M. Evaluation of the Bruker MALDI Biotyper for identification of Gram-positive rods: development of a diagnostic algorithm for the clinical laboratory. J Clin Microbiol. 2014 Apr;52(4):1089– 97.

27. The European Committee on Antimicrobial Susceptibility Testing (EUCAST). When there are no breakpoints in breakpoint tables? EUCAST Guidance; Revised July 2026. https://www.eucast.org/fileadmin/eucast/pdf/guidance_documents/When_there_are_no_breakpoints_revision_20260710.pdf

28. Andrews S. FastQC: A Quality Control Tool for High Throughput Sequence Data. Babraham Bioinformatics; 2010. [cited 2026 Jun 20]. https://www.bioinformatics.babraham.ac.uk/projects/fastqc/

29. Bolger AM, Lohse M, Usadel B. Trimmomatic: a flexible trimmer for Illumina sequence data. Bioinformatics. 2014 Aug 1;30(15):2114–20.

30. Wood DE, Lu J, Langmead B. Improved metagenomic analysis with Kraken 2. Genome Biol. 2019 Nov 28;20(1):257.

31. Seemann T. Shovill [internet]. GitHub; 2016. [cited 2026 Jun 20]. https://github.com/tseemann/shovill

32. Seemann T. Prokka: rapid prokaryotic genome annotation. Bioinformatics. 2014 Jul 15;30(14):2068–9.

33. Page AJ, Cummins CA, Hunt M, Wong VK, Reuter S, Holden MT, et al. Roary: rapid large-scale prokaryote pan genome analysis. Bioinformatics. 2015 Nov 15;31(22):3691–3.

34. Price MN, Dehal PS, Arkin AP. FastTree 2--approximately maximum-likelihood trees for large alignments. PLoS One. 2010 Mar 10;5(3):e9490.

35. Wick RR. Filtlong: quality filtering tool for long reads [internet]. Github. [cited 2026 Jun 20]. https://github.com/rrwick/Filtlong

36. Bouras G, Judd LM, Edwards RA, Vreugde S, Stinear TP, Wick RR. How low can you go? Short-read polishing of Oxford Nanopore bacterial genome assemblies. Microb Genom. 2024 Jun;10(6).

37. Seemann T. ABRicate: Mass screening of contigs for antimicrobial and virulence genes. GitHub. [cited 2026 Jun 20]. https://github.com/tseemann/abricate

38. Jia B, Raphenya AR, Alcock B, Waglechner N, Guo P, Tsang KK, et al. CARD 2017: expansion and model-centric curation of the comprehensive antibiotic resistance database. Nucleic Acids Res. 2017 Jan 4;45(D1):D566–D73.

39. Branford I, Johnson S, Chapwanya A, Zayas S, Boyen F, Mielcarska MB, et al. Comprehensive Molecular Dissection of Dermatophilus congolensis Genome and First Observation of tet(Z) Tetracycline Resistance. Int J Mol Sci. 2021 Jul 1;22(13).

40. Nowicka D, Baglaj-Oleszczuk M, Maj J. Infectious diseases of the skin in contact sports. Adv Clin Exp Med. 2020 Dec;29(12):1491–5.

41. Peterson AR, Nash E, Anderson BJ. Infectious Disease in Contact Sports. Sports Health. 2019 Jan/Feb;11(1):47–58.

42. Poisson DM, Rousseau D, Defo D, Esteve E. Outbreak of tinea corporis gladiatorum, a fungal skin infection due to Trichophyton tonsurans, in a French high level judo team. Euro Surveill. 2005 Sep;10(9):187–90.

43. Balic A, Bukvic Mokos Z, Marinovic B, Ledic Drvar D. Tatami Mats: A Source of Pitted Keratolysis in a Martial Arts Athlete? Acta Dermatovenerol Croat. 2018 Apr;26(1):68–70.

44. McIntyre KM, Setzkorn C, Hepworth PJ, Morand S, Morse AP, Baylis M. Systematic Assessment of the Climate Sensitivity of Important Human and Domestic Animals Pathogens in Europe. Sci Rep. 2017 Aug 2;7(1):7134.

