## Supplementary materials for "Outbreak of *Dermatophilus congolensis* skin infection among contact sport practitioners, Norway, summer 2025"

Contents:

- Figure 4
- Consent form
- Interview guide
- Control measures - details

### Figure 4

Pairwise core-genome SNP distances between the Trondheim *D. congolensis* outbreak isolates (yellow) and international outbreak isolates from Spain (blue) and France (green).

| **Pairwise core genome SNP distances (1,518,281 bp)** | **Range distance to Trondheim strains** | | **Minimum distance to Trondheim strains** | SO-DCO25-1 | SO-DCO25-3 | SO-DCO25-4 | SO-DCO25-5 | SO-DCO25-6 | SO-DCO25-7 | SO-DCO25-8 | SO-DCO26-1 | SO-DCO25-2 |
|---|---|---|---|---|---|---|---|---|---|---|---|---|
| SO-DCO25-1 | | | 0 | | 0 | 0 | 3 | 3 | 3 | 3 | 3 | 4 |
| SO-DCO25-3 | | | 0 | 0 | | 0 | 3 | 3 | 3 | 3 | 3 | 4 |
| SO-DCO25-4 | | | 0 | 0 | 0 | | 3 | 3 | 3 | 3 | 3 | 4 |
| SO-DCO25-5 | | | 0 | 3 | 3 | 3 | | 0 | 0 | 0 | 2 | 7 |
| SO-DCO25-6 | 0 | 7 | 0 | 3 | 3 | 3 | 0 | | 0 | 0 | 2 | 7 |
| SO-DCO25-7 | | | 0 | 3 | 3 | 3 | 0 | 0 | | 0 | 2 | 7 |
| SO-DCO25-8 | | | 0 | 3 | 3 | 3 | 0 | 0 | 0 | | 2 | 7 |
| SO-DCO26-1 | | | 2 | 3 | 3 | 3 | 2 | 2 | 2 | 2 | | 7 |
| SO-DCO25-2 | | | 4 | 4 | 4 | 4 | 7 | 7 | 7 | 7 | 7 | |
| Barcelona_Patient2 | | | 1 | 2 | 2 | 2 | 1 | 1 | 1 | 1 | 1 | 6 |
| Barcelona_Patient6 | | | 1 | 2 | 2 | 2 | 1 | 1 | 1 | 1 | 1 | 6 |
| Barcelona_Patient1 | | | 2 | 3 | 3 | 3 | 2 | 2 | 2 | 2 | 2 | 7 |
| Barcelona_Patient3 | 1 | 7 | 2 | 3 | 3 | 3 | 2 | 2 | 2 | 2 | 2 | 7 |
| Barcelona_Patient4 | | | 2 | 3 | 3 | 3 | 2 | 2 | 2 | 2 | 2 | 7 |
| Barcelona_Patient5 | | | 2 | 3 | 3 | 3 | 2 | 2 | 2 | 2 | 2 | 7 |
| Barcelona_Patient7 | | | 2 | 3 | 3 | 3 | 2 | 2 | 2 | 2 | 2 | 7 |
| Lyon_Patient3 | | | 3 | 4 | 4 | 4 | 3 | 3 | 3 | 3 | 3 | 8 |
| Lyon_Patient1 | 3 | 14 | 7 | 8 | 8 | 8 | 7 | 7 | 7 | 7 | 7 | 12 |
| Lyon_Patient7 | | | 9 | 10 | 10 | 10 | 9 | 9 | 9 | 9 | 9 | 14 |
| Lyon_Patient5 | | | 13 | 14 | 14 | 14 | 13 | 13 | 13 | 13 | 13 | 18 |
| Lyon_Patient6 | | | 16 | 17 | 17 | 17 | 16 | 16 | 16 | 16 | 16 | 19 |
| Lyon_Patient4 | 13 | 52 | 24 | 25 | 25 | 25 | 24 | 24 | 24 | 24 | 24 | 29 |
| Lyon_Patient2 | | | 28 | 29 | 29 | 29 | 28 | 28 | 28 | 28 | 28 | 33 |
| Lyon_Patient8 | | | 47 | 48 | 48 | 48 | 47 | 47 | 47 | 47 | 47 | 52 |
| SO-DCO24-1 | 61 614 | 61 677 | 61 614 | 61 663 | 61 667 | 61 672 | 61 662 | 61 614 | 61 634 | 61 666 | 61 677 | 61 671 |

### Consent form - publication of case in a journal

This is a request for your permission to use information about you and your health for publication in a scientific journal. The condition for which you received treatment is of interest to others working in healthcare. We therefore request your permission to describe this case in articles that may be published in Norwegian or international medical journals.

You will not be directly identifiable in the published material. Your name, personal identification number, or other direct identifiers will not be included. However, it is possible that people familiar with your history may recognize that the information relates to you and, in that way, know more about you and your health.

The information we request permission to use includes data collected by the hospital, your general practitioner, and the municipal communicable disease control service in connection with the treatment you received. Specifically, we wish to use information regarding the infection, any photographs of wounds or skin lesions/rashes, the mode of transmission, treatment, and progression of the disease, as well as general demographic information at the group level (such as sex and age). If you wish, you may review the material once it is completed and before it is used.

**Possible benefits and disadvantages**
You will not have any direct benefit from providing consent. However, publication of experiences from your case may be helpful for healthcare professionals elsewhere.

**Voluntary participation and access to information**
If you do not want us to use information about you in this way, you do not need to provide a reason, and your decision will have no consequences for any future treatment or care you receive.

If you agree, you will sign the consent declaration. Published articles will be available both online and in print. If you wish, you may review the article before publication. Once the article has been submitted to the journal, it will no longer be possible to withdraw your consent.

You have the right to access the information stored about you and to request correction of any inaccurate information.

**Contact information**
If you would like more information, or if you later wish to withdraw, please contact:
Sveinung Javnes
Resident Physician
Department of Medical Microbiology, St. Olavs University Hospital

#### Consent

I consent to information about me and my health being used in articles, as described in the information above.

---------------------------------------------------------------------------------------------------------------
(Patient signature, date)

#### Confirmation that information has been provided to the patient

I confirm that I have provided the information described above

---------------------------------------------------------------------------------------------------------------
(Signature, position/title, date)

### Interview Guide – Skin Infection with Dermatophilus congolensis

Name of interviewer: ..................................................................... Date of interview: ...........................................

**Introduction: Purpose and Confidentiality**

*This interview is conducted as part of the contact tracing and outbreak investigation that health services are required to carry out during outbreaks of infectious diseases. The purpose is to identify possible sources of infection and understand the course of illness, so that we can improve prevention and management of similar outbreaks in the future. The bacterium identified in this case has never previously been described as a cause of a human outbreak in Norway. Therefore, the knowledge gained from this outbreak is of great value.*

*By participating in this interview, you consent to Trondheim Municipality Vaccination and Communicable Disease Control Office, the Microbiology Laboratory at St. Olavs Hospital, and the Norwegian Institute of Public Health using the information you provide to investigate the outbreak. The interviewer is a healthcare professional bound by confidentiality, and your information will be treated confidentially. After the interview, we will ask whether your information may also be used anonymously in a scientific publication. This is voluntary and requires separate consent from you. More information about this will be provided after the interview itself.*

#### ■ Information about the participant

| |
|---|
| Name: |
| Age: |
| Address: |
| Occupation: |
| Workplace: |
| Who do you live with? |
| Telephone number (for future contact): |

#### ■ Information about the skin infection and treatment

| |
|---|
| When did you first notice symptoms of the skin infection? |
| How long did it last? |
| Are you completely recovered/symptom-free now? |

**What symptoms did you have?**

| | Yes | No | Unsure | |
|---|---|---|---|---|
| Rash | ☐ | ☐ | ☐ | ***If yes:***<br>When did it start?<br>What did it look like?<br>Where on the body was it located?<br>How long did it last? |
| Itching | ☐ | ☐ | ☐ | ***If yes:***<br>When did it start?<br>How long did it last? |
| Fever | ☐ | ☐ | ☐ | ***If yes:***<br>When did it start?<br>How long did it last? |
| Other symptoms? | ☐ | ☐ | ☐ | ***If yes:***<br>What symptom(s)?<br>When did it start?<br>How long did it last? |

| |
|---|
| Which symptoms appeared first, which were the most prominent, and how did the infection develop over time? |
| Did you have any wounds or skin injuries in the area where the rash first appeared? After shaving? |

**Medical consultation, treatment and infection control measures**

| | Yes | No | Unsure | |
|---|---|---|---|---|
| Did you contact a doctor? | ☐ | ☐ | ☐ | ***If yes:***<br>When?<br>Which doctor/healthcare provider? |
| Was a sample taken from the rash? | ☐ | ☐ | ☐ | |
| Did you receive topical treatment (washing solution or cream/ointment)? | ☐ | ☐ | ☐ | ***If yes:***<br>What treatment did you receive (name)?<br>When?<br>Was it effective? |
| Did you receive oral antibiotic treatment? | ☐ | ☐ | ☐ | ***If yes:***<br>What antibiotic did you receive (name)?<br>When?<br>Was it effective? |
| Have you taken any special precautions when washing training clothes/equipment, bed linen, or underwear? | ☐ | ☐ | ☐ | ***If yes:***<br>At what temperature were the clothes washed?<br>What measures were taken? |
| Have cleaning or other infection-control measures been implemented where you train? | ☐ | ☐ | ☐ | ***If yes:***<br>What was done?<br>Mats? Other contact surfaces? |

#### ■ Information about other ill individuals

**Are you aware of any other people who have or have had the same symptoms as you?**

| | |
|---|---|
| Yes ☐ No ☐ Unsure ☐ | ***If yes***, how many people? |

| ***If yes*** | Yes | No |
|---|---|---|
| Do these individuals belong to the same household as you? | ☐ | ☐ |
| ***If yes:*** | | |
| • What is your relationship to them? | | |
| • Did they develop symptoms before or after you? | | |
| • How did their skin infection present? | | |
| Do these individuals belong to the same training environment as you? | ☐ | ☐ |
| ***If yes:*** | | |
| • Which training environment? | | |
| • Did you train together (same session/class)? | ☐ | ☐ |
| • Did you have skin-to-skin contact? | ☐ | ☐ |
| • Did they develop symptoms before or after you? | | |
| Do these individuals belong to other settings that you attend? | ☐ | ☐ |
| ***If yes:*** | | |
| • Which setting(s)? | | |

| |
|---|
| • Describe the type of relationship/contact |

#### ■ Information about possible source of infection

| **What do you think caused the illness? (Do you suspect a particular event? Why?)** |
|---|

| **Contact sports** | Yes | No |
|---|---|---|
| Do you participate in contact sports or martial arts? | ☐ | ☐ |
| ***If yes*** | | |
| • Which type(s)? (one or more) | | |
| • Where did you train during the 3–4 months before symptoms started? (Name, location) | Name, place | |
| • Did you train or compete in other cities or abroad during the 3–4 months before symptoms started? (Name, location) | Name, place | |
| • How many training sessions per week did you attend during the 3–4 months before symptoms started | | |
| • What type of clothing did you wear during training? | | |

| **Animal contact** | Yes | No |
|---|---|---|
| During the 3–4 months before your symptoms started, did you have close contact with animals (especially horses, cattle, goats, or sheep) ***in Norway***? | ☐ | ☐ |
| ***If yes:*** | | |
| • Which animals, where and when? | | |
| During the 3–4 months before your symptoms started, did you have close contact with animals (especially horses, cattle, goats, or sheep) ***abroad***? | ☐ | ☐ |
| ***If yes:*** | | |
| • Which animals, where and when? | | |
| Has anyone you live with had close contact with animals during the last 3–4 months (especially horses, cattle, goats, or sheep), either in Norway or abroad? | ☐ | ☐ |
| ***If yes:*** | | |
| • Which animals, where and when? | | |

| **Travel and swimming** | Yes | No |
|---|---|---|
| Did you stay abroad during the 3–4 months before your symptoms began? | ☐ | ☐ |
| ***If yes:*** | | |
| • Where and when? | | |
| Had any of the people you live with travelled abroad during the month before your symptoms began? | ☐ | ☐ |
| ***If yes:*** | | |
| • Where and when? | | |
| Did you travel within Norway during the month before your symptoms began? | ☐ | ☐ |

| | | |
|---|---|---|
| ***If yes:*** | | |
| • Where and when? | | |
| Did you swim in freshwater during the month before your symptoms began? | ☐ | ☐ |
| ***If yes:*** | | |
| • Where and when? | | |

#### ■ Contact information and consent

Email address: ___________________________________

We plan to write a scientific article about this outbreak in order to share the knowledge gained. It would be very useful to be able to use the information you have provided in this interview. If you are willing, we can send you additional information and a consent form by email, which you may sign and return to us.

**Thanks for the information!**

#### Detailed Timeline of Outbreak Investigation and Control Measures

| Date | Action / Control Measures |
|---|---|
| 6 August | *Two initial cases (Cases 1–2): One case reported by a GP, at the same time information from laboratory about two confirmed cases*.<br>The Municipal Communicable Disease Control Service initiated an outbreak investigation. Communication was established with the Microbiology Laboratory at St. Olavs Hospital to verify findings, discuss diagnostic methods, and coordinate further testing. The leadership of the affected sports club was informed and involved in the response.<br>The Norwegian Institute of Public Health (NIPH) was notified and consulted regarding case management, contact tracing, and outbreak control. |
| 7 August | Initial control recommendations were developed and communicated. These included active case finding, testing of symptomatic individuals, temporary exclusion from contact training for suspected cases, reinforcement of personal hygiene measures, recommendations against sharing towels, clothing, or personal equipment, and increased attention to skin lesions among athletes. |
| 8 August | Written information was distributed to all members of the sports club. The information described the symptoms of infection, advised athletes with skin lesions to seek medical assessment and microbiological testing, recommended temporary abstinence from training until recovery, and advised daily body washing with chlorhexidine gluconate (Hibiscrub®). Members were also reminded to wash training clothing and equipment regularly.<br><br>August 8, 2025<br>**Information on Skin Infection Caused by Dermatophilus congolensis**<br><br>Several cases of skin infection caused by Dermatophilus congolensis have been identified among individuals associated with [redacted]<br><br>The infection is caused by the bacterium Dermatophilus congolensis. This is a rare infection in humans, and human infection typically occurs following contact with infected animals in tropical or subtropical regions.<br><br>Transmission usually occurs through direct contact, particularly when the skin is damaged or has open wounds.<br><br>The infection typically begins with scattered itchy spots on the skin, which gradually develop into oozing and crusted lesions. The rash may spread to multiple areas of the body. Fever may also occur.<br><br>The bacterium thrives in wet and humid conditions, which may occur in training facilities of this type, and it can survive in the environment for a period of time.<br>Good hygiene and local wound care are important. Showering with Hibiscrub soap every other day is recommended until the infection has healed.<br><br>Individuals with skin infection should not participate in training activities until complete recovery.<br>If the skin infection spreads to several parts of the body, a general practitioner should be consulted to assess the need for treatment. Antibiotic therapy may be required if the infection is extensive and does not improve with local treatment alone.<br><br>Measures taken at the training facility include thorough cleaning of the premises, contact surfaces, training mats, and other equipment using soap and water. |

<table>
<tr><td></td><td>
For questions, please contact the Vaccination and Infection Control Office, Trondheim Municipality, at +….. (daytime hours).<br><br>
Sincerely,<br>
Trondheim Municipality<br>
Municipal Medical Officer for Communicable Disease Control
</td></tr>
<tr><td>11 August</td><td>
Written outbreak information was sent to all general practitioners and the local out-of-hours medical service. The notice described the clinical presentation, raised awareness of the outbreak, provided guidance on sampling and microbiological diagnostics, and encouraged healthcare professionals to consider Dermatophilus congolensis in patients presenting with compatible skin lesions.<br><br>
August 11, 2025<br>
Dermatophilus congolensis Identified in Skin Infections Among Members of a Sports Club<br>
Information for General Practitioners and Out-of-Hours Physicians<br><br>
Skin infections caused by Dermatophilus congolensis have been identified in several persons training at [redacted]. This is a rare infection in humans and typically occurs following contact with infected animals in tropical or subtropical regions.<br><br>
Transmission usually occurs through direct contact, particularly when the skin is damaged or has open wounds.<br><br>
The infection typically begins as scattered itchy spots that gradually develop into oozing and crusted lesions. The rash may spread to multiple body sites. Fever may also occur.<br><br>
The bacterium thrives under wet and humid conditions, as may occur in training facilities, and can survive in the environment for a period of time.<br><br>
Good hygiene and local wound care are important. Showering with Hibiscrub soap every other day is recommended until the infection has resolved. Individuals with skin infection should not attend training until fully healed.<br><br>
When skin infection spreads to multiple parts of the body, patients should contact their GP for assessment. Antibiotic treatment may be necessary if the infection is extensive and does not improve with local treatment.<br><br>
If infection with this bacterium is suspected, microbiological samples should be collected from the skin lesions. The laboratory request form should be marked as suspected outbreak-related D. congolensis infection. Samples should preferably be sent to the Department of Medical Microbiology, St. Olavs Hospital.<br><br>
Measures implemented at the training facility include thorough cleaning of the premises, contact surfaces, training mats, and other training equipment using soap and water.<br>
The members of the sports club have been informed through the club management. This information has been prepared in collaboration with the Department of Dermatology at St. Olavs Hospital and the Norwegian Institute of Public Health.
</td></tr>
<tr><td>13 August</td><td>Report from a GP regarding a new case (Case 4)</td></tr>
<tr><td>18 August</td><td>
Information on a third case (Case 3) via leader of a second sportsclub<br>
Written information was provided to an additional sports club attended by one of the confirmed cases. The club was informed about the ongoing outbreak, advised to increase vigilance for skin symptoms among members, and encouraged to implement preventive hygiene measures and facilitate testing of symptomatic individuals. No further cases were found related to this club.
</td></tr>
</table>

| | |
|---|---|
| 18–25 August | Training facility were temporarily closed to interrupt possible transmission. During the closure period, the premises underwent extensive cleaning and disinfection of mats, training equipment, changing rooms, and high-touch surfaces. Facility was also aired and maintenance routines reviewed before reopening. |
| 22 August | An updated advisory was distributed to all general practitioners and the out-of-hours medical service. Clinicians were requested to report suspected cases directly to the Municipal Chief Medical Officer (CMO) to strengthen surveillance, facilitate rapid case assessment, and improve situational awareness during the outbreak.<br><br>August 22, 2025<br>**Status Update: Outbreak of Skin Infection Caused by Dermatophilus congolensis**<br>Information for General Practitioners and Out-of-Hours Physicians<br><br>We have received notifications of several cases of skin infection caused by Dermatophilus congolensis associated [redacted]. An additional case has also been identified at [redacted]<br><br>It is therefore important to obtain samples from skin infections when infection with this bacterium is suspected. This also applies to persons training at other sports facilities, with particular attention to those participating in martial arts. The laboratory request form should be marked as suspected outbreak-related D. congolensis infection. Samples should preferably be sent to St. Olavs Hospital.<br><br>The literature also describes co-infection with other microorganisms. Ringworm (dermatophyte infection) and Staphylococcus aureus infections are particularly common among practitioners of martial arts such as [redacted].<br>To maintain oversight of the outbreak, please send an electronic message to Trondheim Municipality Infection Prevention and Control Services whenever Dermatophilus congolensis is identified.<br><br>Important reminders from the information distributed on August 11, 2025<br>• Good hygiene and local wound care remain essential. Showering with Hibiscrub soap every other day is recommended until the infection has healed.<br>• Antibiotic treatment may be necessary if the infection is extensive and does not improve with local treatment.<br>• Persons with skin infection should not participate in training sessions or competitions until the infection has healed. This also applies to training at other facilities.<br>• Training facilities should conduct thorough cleaning of premises, contact surfaces, training mats, and equipment using soap and water. Training clothes and uniforms should be washed at 60°C (140°F). |
| 25 August | *Four additional cases (Cases 5–8) were confirmed.* Epidemiological investigations were intensified, including further contact tracing, interviews of affected individuals, mapping of possible transmission routes, and assessment of additional control measures. |
| 25–26 August | The outbreak received attention in local and subsequently national media following media inquiries. Although media communication was not originally planned as part of the outbreak response strategy, the CMO provided factual information about the outbreak, symptoms, and ongoing investigations. |
| 17 September | *One additional case (Case 9) was confirmed* despite the implemented control measures. Continued surveillance, follow-up of affected individuals, and monitoring for new cases were maintained throughout the investigation period. |
